# Predicting transcriptional responses to cold stress across plant species

**DOI:** 10.1101/2020.08.25.266635

**Authors:** Xiaoxi Meng, Zhikai Liang, Xiuru Dai, Yang Zhang, Samira Mahboub, Daniel W. Ngu, Rebecca L. Roston, James C. Schnable

**Author notes:** Department of Horticultural Science, University of Minnesota, St. Paul, MN 55108 USA. Department of Plant and Microbial Biology, University of Minnesota, St. Paul, MN 55108 USA. St. Jude Children’s Research Hospital, Memphis, TN 38105 USA.

## Abstract

Although genome sequence assemblies are available for a growing number of plant species, gene expression responses to stimuli have been catalogued for only a subset of these species. Many genes show altered transcription patterns in response to abiotic stresses. However, orthologous genes in related species often exhibit different responses to a given stress. Accordingly, data on the regulation of gene expression in one species are not reliable predictors of orthologous gene responses in a related species. Here, we trained a supervised classification algorithm to identify genes that transcriptionally respond to cold stress. A model trained with only features calculated directly from genome assemblies exhibited only modest decreases in performance relative to models trained using genomic, chromatin, and evolution/diversity features. Models trained with data from one species successfully predicted which genes would respond to cold stress in other related species. Cross-species predictions remained accurate when training was performed in cold-sensitive species and predictions were performed in cold-tolerant species and vice versa. Models trained with data on gene expression in multiple species outperformed models trained with data from any single species. These results suggest that classifiers trained on stress data from well-studied species may suffice for predicting gene expression patterns in related, less-studied species with sequenced genomes.

## Introduction

The genomes of over 300 plant species have been sequenced to date. Ambitious efforts are underway to sequence the genomes of up to 10,000 plant and algae species by 2023^1^. Even members of closely related groups of species can be adapted to different environments and exhibit different degrees of tolerance for different stresses. The panicoid grasses are a clade of approximately 3,000 plant species, including several domesticated crops. While panicoid grasses grow in and are adapted to a wide range of environments, many of the most agriculturally and economically important species, including maize (*Zea mays* ssp. *mays*) and sorghum (*Sorghum bicolor*), were originally domesticated at tropical latitudes and are not cold tolerant. For these crops, the low-temperatures in the spring and autumn constrain the length of the growing season and pose a major limit to total agricultural production. While the majority of panicoid grasses are native to the tropics or subtropics^2^, a number of lineages have evolved to grow in temperate environments where cold and freezing temperatures occur annually. For instance, miscanthus (*Miscanthus giganteus*), a cold-tolerant relative of maize and sorghum that is native to temperate environments, exhibits substantially higher total photosynthetic productivity per year than these crops due to its longer growing season and reduced susceptibility to photoinhibition at chilling temperatures1^3^. Thus, the clade contains a complex mixture of cold-tolerant species such as foxtail millet (*Setaria italica*) and switchgrass (*Panicum virgatum*)^4,5^ and cold-sensitive species including maize, sorghum (*Sorghum bicolor*), proso millet (*Panicum miliaceum*), and pearl millet (*Pennisetum glaucum*).

Plants have evolved a variety of physiological, biochemical, and transcriptional regulatory mechanisms to sense and respond to abiotic stress^6^. The repeated acquisition and/or loss of cold tolerance within the panicoid grasses provides an opportunity to better understand the biochemical and evolutionary mechanisms responsible for changes in temperature tolerance. However, the patterns of gene expression variation in response to cold stress are not conserved across species^7,8^ or even between genotypes within the same species^9^. The modulation of transcriptional regulation in response to abiotic stress often requires synchronous actions among cis-regulatory elements (e.g. promoter and enhancer), trans-regulatory elements (e.g. transcription factor and regulating RNA), transposable elements, and epigenetic regulators (e.g. DNA methylation and chromatin structure)^6,9–11^.

One explanation for the rapid divergence of cold-responsive transcriptional regulation between orthologous genes is that new insertions of transposable elements appear to have the potential to induce the cold-responsive expression of nearby genes^11–13^. It is likely that the rewiring of transcriptional regulation plays a significant role in how different plant lineages adapt independently to low-temperature stress.

Here we demonstrate that even though orthology is not an effective predictor of transcriptional responses to cold stress across even closely related species, it is possible to train supervised classification models using data from one species to predict which genes will respond to cold stress in another species. The usefulness of supervised classification algorithms has been demonstrated for a range of biological applications, such as distinguishing gene models with the potential for expression^14^, inferring human gene expression based on a mouse model^15^, predicting functional annotations of individual gene models from functional genomic data^16^, distinguishing genes involved in specialized or primary metabolism^17^, and predicting post-translational modification sites^18^. In this study, we generated transcriptional data from four closely related species: foxtail millet, pearl millet, switchgrass, and proso millet (Figure1A). Importantly, models used to predict which genes would transcriptionally respond to cold stress provided equivalent prediction accuracy when trained using only features calculated from the genome and gene model annotations as when trained using larger feature sets that included evolutionary, chromatin, and population diversity features. With genome-and-gene-model-only feature sets, models trained in one species could be used to make predictions in a second species. This cross-species prediction method provides an effective mean of predicting which genes will transcriptionally respond to cold stress without the need to generate new expression datasets under equivalent conditions for each species. With the growing number of sequenced plant genomes, the ability to predict transcriptionally responded genes to stresses based on data from genome sequence assemblies will lower the barriers to investigating the basis of widespread variation in stress tolerance across the plant kingdom.

**Figure 1.**
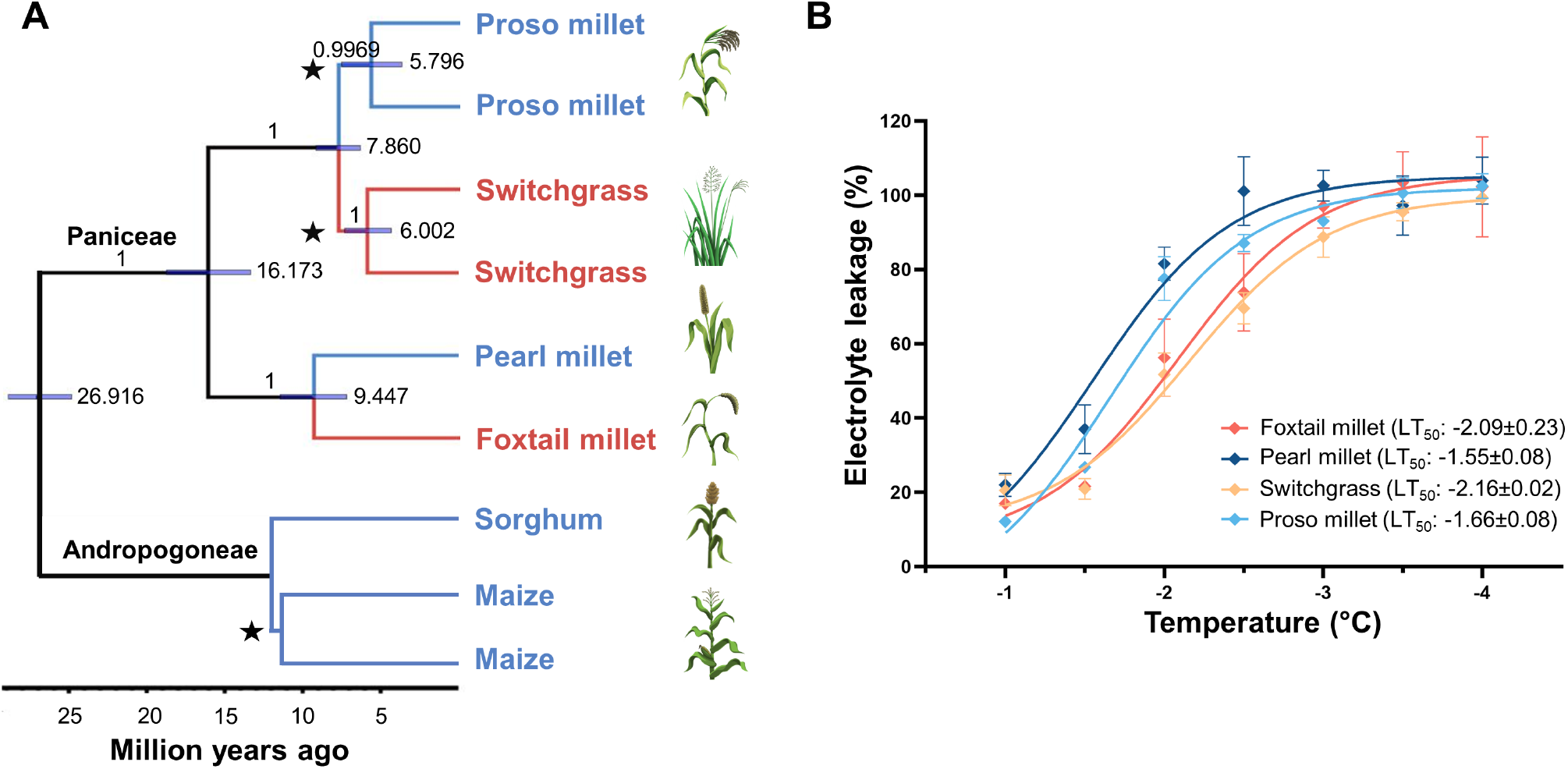
Phylogenetic and phenotypic relationships between foxtail millet, pearl millet, switchgrass, and proso millet. A. Species tree for the six species investigated in this study. Branches shown in red are relatively cold tolerant compared to branches shown in blue. Branch supports are Bayesian posterior probabilities, node bars show 95% highest posterior density (95% HPD) of node age and the scale bar represents millions of years ago. The whole genome duplication events are marked by stars and indicating that the species contains two subgenomes. Maize was not included during species tree analysis and the divergence time between maize and sorghum was calibrated to 11.9 million years ago^22^. B. Electrolyte leakage from non-acclimated leaves frozen to a range of different temperatures. Curves were fitted using nonlinear regression with a sigmoidal dose-response model. LT50 values are the concentrations that give half-maximal effects^23^. Error bars indicate standard error of the mean from at least three replicate measurements.

## Results

### Cold-Responsive Genes and Gene Expression Patterns Vary Among Related Species

Both maize and sorghum are sensitive to cold stress^4,7,19,20^. Reports of the differences in the degrees of low-temperature tolerance among Paniceae species are sparse and varied, although switchgrass is extremely tolerant of cold and freezing, at least under some conditions^4,5,21^. Cold tolerance can vary substantially depending on treatment, developmental stage, and acclimation (as previously reviewed4). Here, we grew seedlings of four Paniceae species under controlled conditions and assayed freezing tolerance at the three-leaf stage using an *in vitro* electrolyte leakage assay resulting from cell breakage to quantify the extent of damage. When not previously acclimated to stress conditions, switchgrass and foxtail millet seedlings showed slower rates of electrolyte leakage when challenged with progressively greater freezing stress compared to pearl millet and proso millet seedlings grown and tested under the same conditions (Figure 1B). Therefore, low-temperature tolerance is not monophyletic within the Paniceae and could reflect the parallel adaptation of different lineages within the grass tribe to temperate climates (Figure 1A and B).

Changes in gene expression induced by cold stress were assayed using paired control and stress treatment RNA-seq datasets collected from foxtail millet, pearl millet, switchgrass, and proso millet at the three-leaf stage 0.5, 1, 3, 6, 16, and 24 hours after the onset of cold stress. Among genes showing cold-responsive changes in mRNA abundance, at least 47% were not syntenically conserved among the four species (Supplementary Table S1 and Figure S1A). The number of nonsyntenic genes that responded transcriptionally to cold stress was more variable across species compared to syntenic genes (Figure S1A). Syntenic orthologous genes and promoters are derived from a single common ancestral gene and promoter of the most recent common ancestor of the species being studied. However, despite this shared evolutionary history, a gene responding transcriptionally to cold stress in one species was not a good predictor of whether syntenic orthologous genes in related species would also respond to cold stress in the same treatment at the same developmental stage (Figure S1B). This low conservation of transcriptional responses across conserved genes in related species is consistent with the results of a previous comparison of the transcriptional responses of maize and sorghum to cold stress7 and the variation in transcriptional responses to cold stress between different alleles of the same gene in maize^9^.

### Supervised Classification Algorithms Can Accurately Predict Cold Responsiveness

Stress-responsive transcriptional regulation of a given gene cannot be predicted efficiently using data from orthologous genes in related species. However, perhaps specific features or properties of the gene itself which can be used to predict whether its expression will respond to cold stress. We first evaluated this approach in maize, as many different types of feature data are available for all or nearly all gene models in this plant^16^. One potential factor that could confound efforts to predict differential gene expression is that the average gene expression level itself is a reasonably good predictor of whether or not a gene will be identified as showing statistically significant differential expression. In the current study, the areas under the receiver operating characteristic curves (AUC-ROCs) for predicting differential expression solely based on average expression levels varied from 0.48 to 0.70 for the six species tested (Figures 2A and S2A). Average gene expression levels can be predicted reasonably well based on genomic features^24^. Clearly, if not controlled for, the association between average gene expression and the odds of a gene being identified as differentially expressed would lead to a misleading estimate of prediction accuracy. We therefore used a gene binning strategy where genes were divided into 12 bins (dodeciles) based on average expression levels and subsampled to ensure equal representation of cold-response and cold nonresponsive genes within each dodecile (Figure 2A, S2B, and 3). After binning and subsampling, the prediction of which genes would be differentially expressed based solely on average expression values produced AUC-ROCs of approximately 0.5, i.e., equal to the null expectation for balanced data (Figures 2A and S2A). We also performed gene-family guided splitting, a strategy proposed by Washburn *et al.*^24^, to avoid obtaining misleadingly high accuracy values that can result when prediction models learn gene family-specific features (Figure 3).

**Figure 2.**
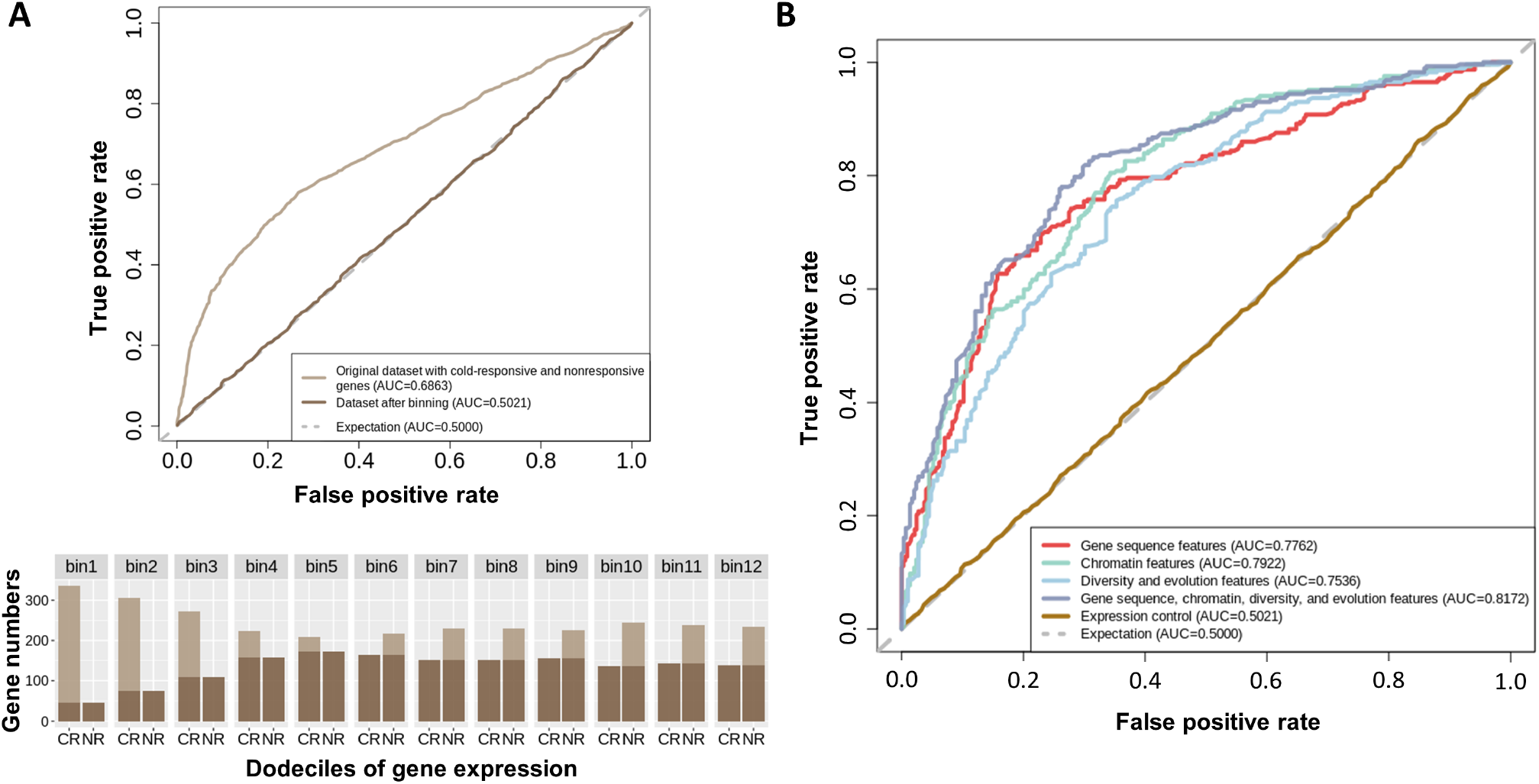
Cold-responsive gene prediction in maize. A. Baseline expression control. Upper: Accuracy of genes being scored as cold-responsive genes solely based on average FPKM values before and after baseline expression control; Lower: Distribution of average FPKM values of cold-responsive genes (CR) and nonresponsive genes (NR), and training sets resampled from genes in 12 bins with balanced gene expression levels (darker color). B. Receiver operating characteristic (ROC) curves showing the classification accuracy of the maize models using different types of features

**Figure 3.**
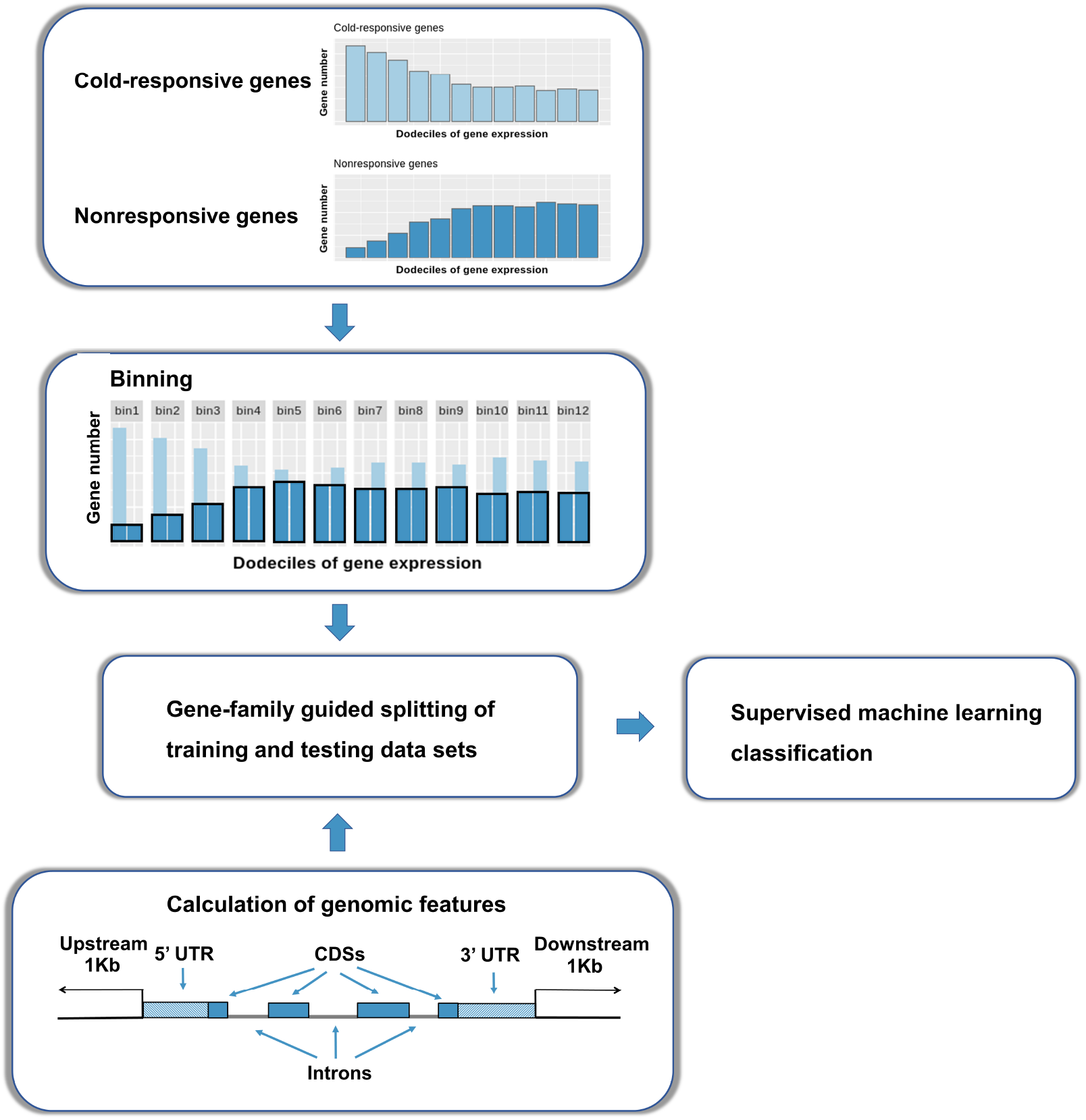
Workflow of the supervised machine classification model for predicting cold-responsive genes.

A set of features was assembled for each maize gene including gene sequence features, chromatin features, and divesity/evolutionary features16 (Supplementary Table S2). Random forest models25 trained using the complete set of features were able to predict which genes would exhibit differential expression in response to cold stress and which would not with an AUC of 0.82 (Figure 2B). Models trained with subsets of features did not match the accuracy of the combined model. A model trained using only features that can be extracted from genomic sequence data was able to predict which genes would exhibit differential expression in response to cold stress and which would not with an AUC of 0.78 (Figure 2B).

Unlike the combined model, which requires data obtained using a range of specialized sequencing techniques, as well as resequencing data from diverse populations, the pure genomic feature model can be applied to any species with a sequenced genome and annotated gene models. We scored the same set of genomic sequence-derived features for each gene model in foxtail millet, pearl millet, switchgrass, and proso millet and trained the species-specific random forest prediction models for each of the four species. The AUC values (indicating the accuracy of species-specific models for classifying genes as cold-responsive or nonresponsive) were 0.83 for foxtail millet, 0.89 for pearl millet, 0.80 for switchgrass, and 0.85 for proso millet (Figure 4A).

**Figure 4.**
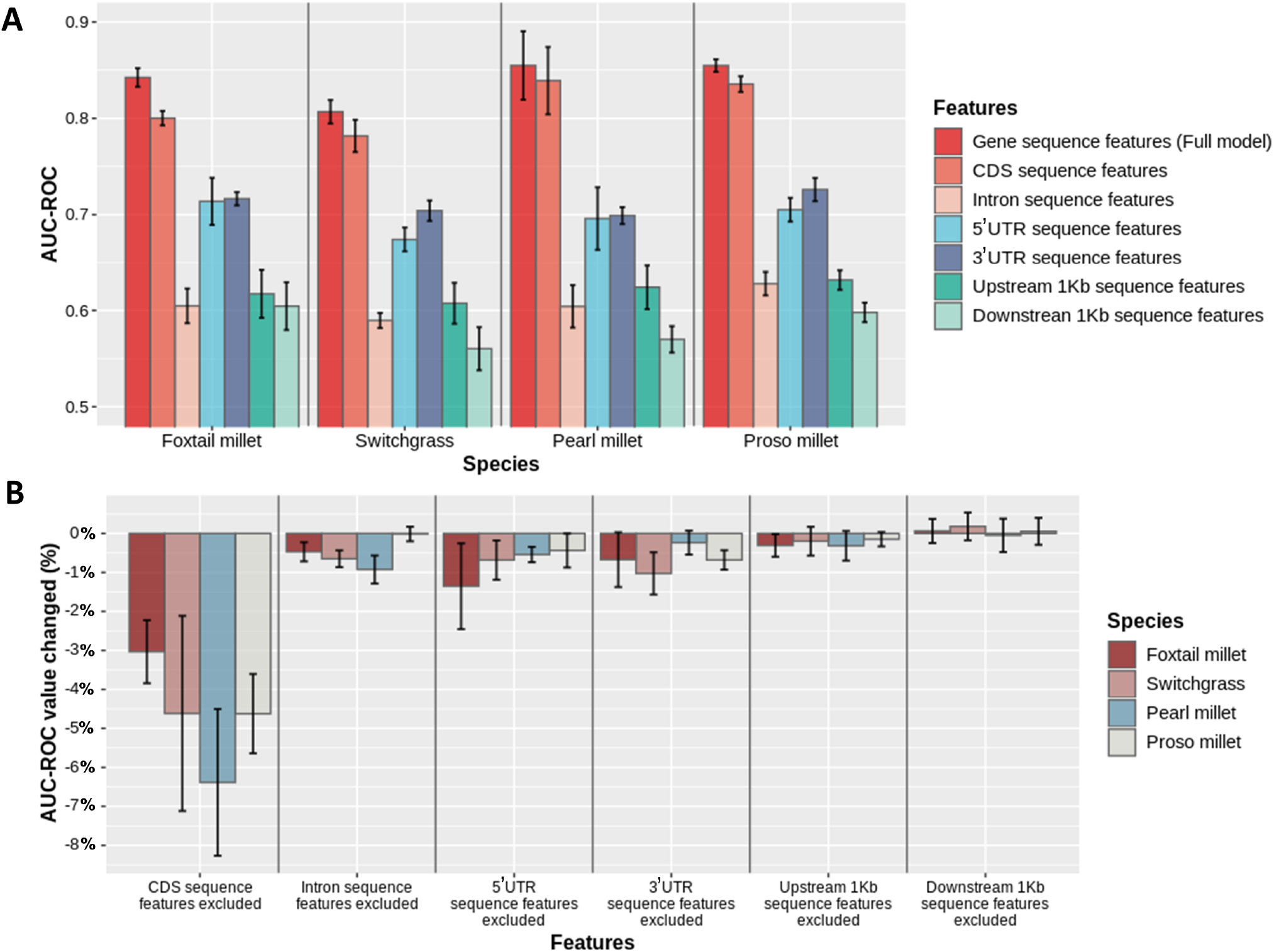
Supervised machine learning models for Paniceae grass species based on gene sequence features. A. Bar plot showing the prediction accuracies achieved by the full gene sequence models and single feature group models for foxtail millet, pearl millet, switchgrass, and proso millet. B. Importance of single feature groups, as revealed by changes in accuracy when leaving one feature group out of the full sequence model. The value (percentage) was calculated based on changes relative to the full model. Negative values indicate a decrease in accuracy compared to the full model, while positive values indicate an increase in accuracy compared to the full model when excluding the feature group.

Each species-specific model showed similar trends in terms of which features were estimated to play the largest roles in determining accuracy. The contents of CG and AA dinucleotides in the CDS region consistently ranked as the two most important features distinguishing cold-responsive genes from genes that did not transcriptionally respond to cold stress. A complete list of estimated importance values for each feature in each species-specific model is provided in Supplementary Table S5. Separate models were trained using only sequence features calculated for specific gene regions: CDS, intron, 5’UTR, 3’UTR, upstream, and downstream regions (Figure 4A). A second set of models was trained using all sequence features except for those calculated for one of the same six regions (Figure 4B). It appears that CDS, 5’UTR, and 3’UTR features provided more useful information for predicting whether or not a given gene model will respond to cold stress than introns, upstream, or downstream regions. CDS-only models consistently performed the best of any of the single sequence context models, but they did not exceed the prediction accuracy of the full model in any of the four species tested.

### Models Trained in One Species Can Predict Cold-Responsive Gene Expression in Another Species

Because the same sequence features can be calculated for genes in different species, it is possible to evaluate how well cold-responsive gene expression can be predicted in one species based on only information about which genes did and did not respond to cold in another species. We used single species models trained in foxtail millet, pearl millet, switchgrass, or proso millet to predict which genes would transcriptionally respond to cold in foxtail millet, pearl millet, switchgrass, proso millet, sorghum, and maize (Figure 5). The accuracy with which cold-responsive gene expression was predicted across species was comparable or only modestly less than the accuracy of within-species prediction. Predictions using species that were more closely related were not obviously, consistently superior to predictions using species that share common cold-stress phenotypes (sensitivity or tolerance).

**Figure 5.**
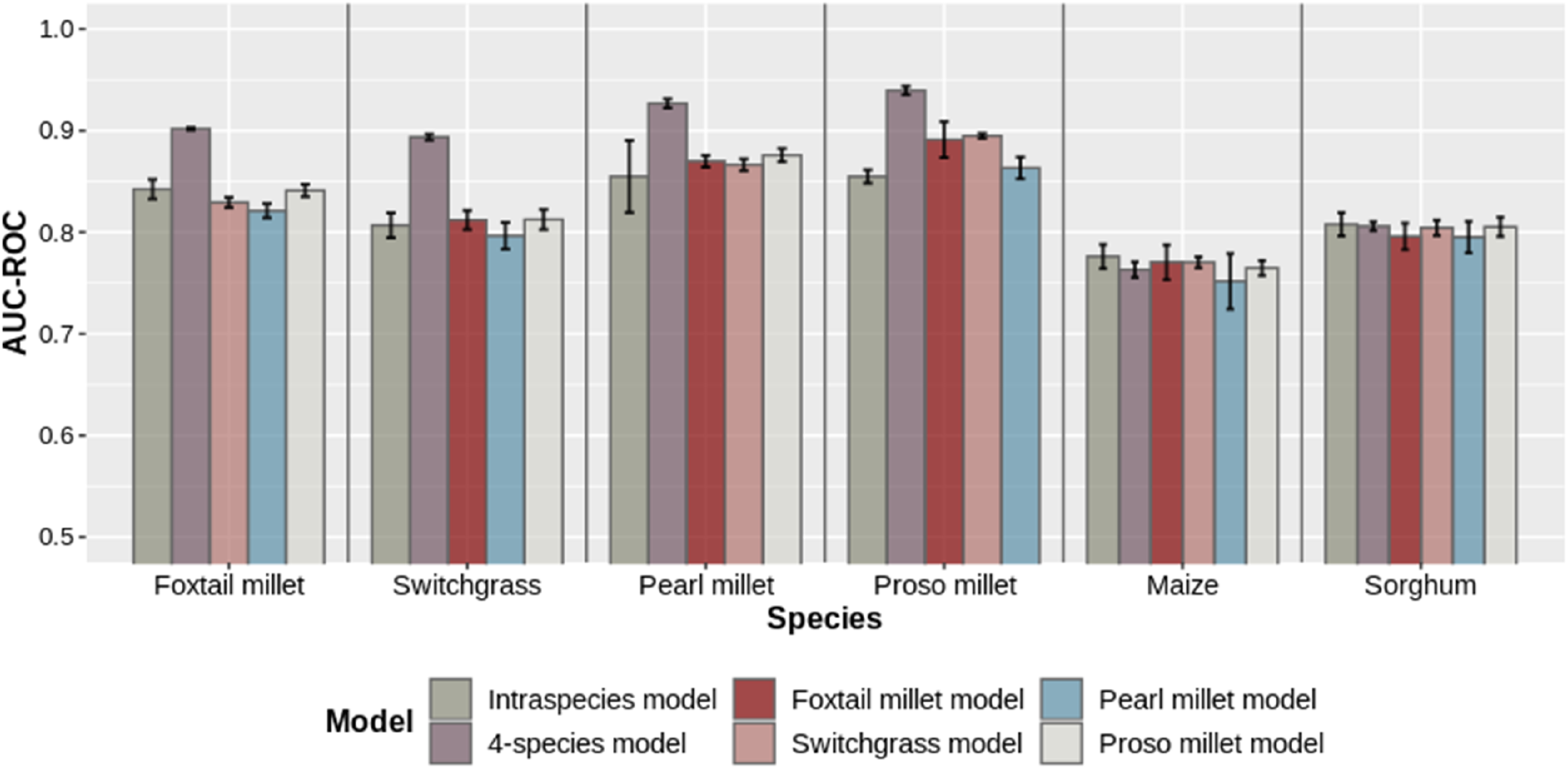
Performance of models trained for cross-species prediction. Prediction accuracy of models trained in foxtail millet, pearl millet, switchgrass, proso millet, or a combination of these four species for classifying cold-responsive genes from nonresponsive genes in Panicoideae grass species including maize, sorghum, and the four species mentioned above. Accuracies of AUC under ROC curves are presented with standard deviations calculated from five-fold cross validation.

A final model was trained using data from all the four Paniceae species (foxtail millet, pearl millet, switchgrass, and proso millet) and its accuracy was assessed using separate data from the same four species as well as data from maize and sorghum. The model trained with data from four species outperformed all cross-species prediction models trained with data from a single species, except in maize. The four species models also outperformed within-species predictions in foxtail millet, proso millet, switchgrass, and sorghum. Unexpectedly, models trained in different species tended to exhibit similar performance when predicting cold-responsive gene expression in the same species, while models trained in a single species tended to exhibit a wider range of performances when predicting cold-responsive gene expression in different species. Models consistently performed the best in classifying either pearl millet or proso millet genes as cold-responsive or nonresponsive and generally performed the worst in predictions from maize. This pattern would be consistent with the notion that a certain proportion of classification errors resulted from varying amounts of noise in the ground truth classifications of gene expression patterns in individual species and/or variation in the accuracy of gene structural annotations used to calculate the sequence features used for prediction.

The successful cross-species predictions for cold-responsive genes between cold-tolerant and cold-sensitive species or between genetically relatively distant species indicate that cold-responsive genes in Panicoideae share a high level of similarity in terms of gene sequence features. The determinants of gene expression under cold stress are consistent across species at the gene sequence level, even though only a small proportion of cold-responsive genes were conserved in different species (Figure S1B). It appears that low-temperature-tolerant species with independent origins employ a similar set of strategies to confer this trait. We further conducted k-mean clustering analysis of cold-responsive genes from the four Paniceae species based on gene expression in response to different durations of cold treatment (Figure S3A-D). Interestingly, we found genes belonging to each of the four species were distributed across all of the clusters (Figure S3E).

## Discussion

Several factors have lowered the barriers to generating reference genome sequences for new species, including declining sequencing costs, advances in long-read sequencing technologies, and improvements to genome assembly and annotation algorithms. To date, over 300 plant genomes have been sequenced; in addition, a recent study collected transcriptome data from 1,124 plant species^26^. Unfortunately, progress in generating layers of functional genomic data, including RNA-seq data for many of these newly sequenced genomes, has been much slower due to issues ranging from seed dormancy and limited access to wild plant species to difficulties in staging plants or delivering controlled stresses, tissues, and cell types, which require complicated, labor-intensive techniques to sample. As mentioned above, methods used to predict stress-responsive gene expression based on data on orthologous genes in related species have low accuracy and would, in any case, likely miss the changes in gene regulation associated with differences in stress tolerance between related species. Instead, we have demonstrated that supervised classification algorithms trained on gene features, including sets of features that can be calculated solely from genomic sequence data and gene structural annotation, can provide significant accuracy to predict which genes will transcriptionally respond to a specific abiotic stress (cold, in this case). The success we achieved in prediction based on gene sequence features greatly expands the potential application of this technique to nonmodel species—including those adapted to extreme environments—for which a reference genome sequence has been generated but substantial functional genomic datasets are lacking.

The application of supervised classification, and machine learning algorithms in general, requires the careful use of controls to avoid getting the right answer for the wrong reasons. Here, we identified three potential risk factors that might produce apparently high accuracy for reasons that would not be biologically meaningful. The data sets used for the supervised machine learning models in our study were well balanced by controlling baseline gene expression between cold-responsive and nonresponsive gene sets and controlling evolutionary relatedness between training and test sets by family-guided training/test splitting. Failure to properly address these factors would likely produce misleading results. Ranking genes by their baseline expression values (average FPKM) allowed us to predict cold-responsive genes with a relatively high performance in most cases (AUCs ranging from 0.48 to 0.70) compared to the expectation by chance (AUC=0.50) (Figure 2A and Supplementary Figure S2), indicating that the accuracy of predicting cold-responsive genes could be confounded by baseline gene expression variation. To accurately evaluate the performance of feature groups and to ensure that genes are classified solely based on their intrinsic features, we controlled the baseline expression bias between cold-responsive and nonresponsive genes using a binning approach after sorting the genes based on average FPKM values. By randomly selecting the same number of genes from cold-responsive and nonresponsive pools in each bin according to the smaller sample size in either pool, the newly constructed data set was balanced, as it contained the same number of cold-responsive genes and nonresponsive genes with comparable baseline expression profiles. After controlling for baseline gene expression, the prediction accuracy for classifying transcriptional responses to cold stress approached the accuracy of expectation by chance (AUC=0.50), implying that our data set for model training was optimized without interference from baseline expression bias. Furthermore, we used the “gene-family guided splitting” method to properly structure the model training and testing data sets to avoid dependencies between the sets. This method ensured that genes within the same family were not split between the training and testing data sets. Genes within or across species could share evolutionary histories, such as gene duplication and gene family relatedness, which would interfere with the model learning only from the desired features. By contrast, simply allocating the genes randomly into training and testing sets without considering their evolutionary relatedness causes model overfitting and can lead to false positive, spurious conclusions^24^. In addition, many stresses are difficult to replicate effectively across different species or laboratories^4^; here, we were able to use plants grown in the same laboratory with consistent growth conditions and treatments, and at the same developmental stage. This consistency represents an advantage that undoubtedly contributed to the success of prediction.

Separate predictions using distinct or ‘leaving-one-out’ subsets revealed that features calculated from CDS, 5’UTR and 3’UTR regions were much more important for obtaining accurate predictions than features calculated from intron, upstream, or downstream regions (Figure 4). The involvement of UTRs in transcriptional regulation was also observed in a study utilizing deep learning^24^, but CDS regions were not analyzed in that study due to technical limitations. In models trained using features calculated from all gene regions, CG dinucleotide content ranked top two in terms of feature importance in foxtail millet, pearl millet, switchgrass, and proso millet. The cytosines in CG sites are active targets of methylation and can be involved in regulating gene expression^27^, but there is also evidence that CG sites can contribute to the regulation of transcriptional activity independently of DNA methylation^28^.

A strikingly low level of conservation of cold responsiveness was observed among syntenic orthologous genes across the species examined in this study (Supplementary Figure S1). The divergence of transcriptional patterns between orthologous genes can result from either trans-regulatory changes or cis-regulatory changes. In a comparison of natural maize haplotypes, cis-regulatory divergence was observed much more frequently than trans-regulatory divergence^9^. The notion that the high degree of divergence in cold-responsive expression between orthologous genes in related species is indeed primarily due to cis-regulatory changes is consistent with the observation that feature importance was conserved between models trained in different species. Specifically, a median of 70% of the 20 features with the highest importance scores overlapped between models trained in different species. In addition, the importance of cis-regulatory changes could explain why the models trained in one species that were successful at predicting cold-responsive gene expression tended to be successful in a second species. However, it is too early to conclude with certainty that features consistently ranked as highly important in multiple models play a causal role in determining whether a gene will transcriptionally respond to cold stress or whether they are simply correlated with this response.

## Materials and Methods

### Plant Material, Growth, and Stress Conditions

For three of the six species tested, we employed the same genotype that had been sequenced to assemble the reference genome for that species: maize (*Zea mays* ssp. *mays* genotype B73), sorghum (*Sorghum bicolor* genotype BTx623), and foxtail millet (*Setaria italica* genotype Yugu1). For the three other species, we were unable to employ the reference genotype and used another variety instead: switchgrass (*Panicum virgatum* genotype kanlow), proso millet (*Panicum miliaceum* genotype earlybird USDA PI 578073), and pearl millet (*Pennisetum glaucum* syn *Cenchrus americanus* genotype USDA PI 583800). For maize and sorghum, gene expression data and details about growth conditions and stress treatments were described in Zhang *et al*. (2017)^7^. Seeds were planted in standard potting mix (40% Canadian peat, 40% coarse vermiculite, 15% masonry sand, and 5% screened topsoil) in a Percival growth chamber (Percival model E-41L2) under 111 mol m^-2^ s^-1^ light intensity, 60% relative humidity, and a 12 h/12 h day-night cycle at 29°C during the day and 23°C at night. To target the approximately three-leaf stage in the different species, planting dates were staggered to allow cold-stress treatments to be performed simultaneously for batches of seedlings from multiple species: foxtail millet, pearl millet, switchgrass, and proso millet seedlings were subjected to cold-stress treatment at 12, 10, 17, and 14 days after planting, respectively. Seedlings at the desired growth stage were divided, with one half of each variety transferred to a growth chamber maintained at 6°C and the other half used as the control. The seedlings were always transferred to cold-stress treatment at the end of the 12-hour day cycle. Paired samples were collected from control and cold-stress treatments at 0.5, 1, 3, 6, 16, and 24 h after the onset of cold stress. Each sample was a pool of all above ground tissue from at least three individual seedlings. Samples were collected from three independent biological replicates grown and harvested on separate dates.

### Electrolyte Leakage Analysis

Plants used for electrolyte leakage analysis were harvested from non-acclimated plants at the same growth stage and conditions described above. Leaf tissue was harvested from pearl millet and proso millet using a 5 mm punch, and three punches were tested per sample. For switchgrass and foxtail millet, the narrow leaf blades prevented the even application of the 5 mm punch; instead, six 5 mm leaf sections were cut with a razor blade and pooled for each sample. Efforts were made to ensure that equivalent portions of the leaf were included in each replicate, and only the midsection of each leaf was used, avoiding the stalk or tip. All leaf samples were immersed in sterile water with a resistivity of 18.2 MΩ at 25°C. All conductivity measurements were performed with an Accumet 200 conductivity meter (Fisher Scientific, probe: catalog number 13-620-101). Initial readings were collected from samples incubated at 0°C for 30 min in a pre-cooled chiller (initial measurement). After pre-chilling, a small ice crystal was added to each sample to initiate ice nucleation. After nucleation, individual samples were incubated in the chiller at a rate of −0.5°C per 0.5 hour, and samples were removed when the temperature reached −1°C, −1.5°C, −2°C, −2.5°C, −3°C, −3.5°C, and −4°C. The samples were thawed at 4°C in a cooling water bath for 2-4 hours, incubated at room temperature for 30 minutes, and mixed on an orbital shaker at 250–300 rpm (rotations per minute) for an additional 20 minutes at room temperature. At this point, the conductivity of the water was measured (treatment measurement). Finally, each sample was incubated at 65°C for 30 minutes and shaken for 20 minutes before a final conductivity reading was taken for each sample (final measurement). Percent electrolyte leakage for each sample was calculated using the formula (treatment measurement - initial measurement)/(final measurement - initial measurement). The LT50 (temperature of 50% electrolyte leakage) for each set of samples was defined to be the LOGEC50 value for a sigmoidal curve fit to the percent leakage values calculated at different temperatures, based on the initial and final measurements. Curves were fit to percent electrolyte leakage value points using the sigmoidal dose-response model provided by the software package GraphPad Prism (v 8.1.2) following the protocol outlined by Thalhammer and coworkers^23^.

### Generating RNA-Seq Data and Identifying Cold-responsive Genes

RNA isolation and library construction were performed as described by Zhang *et al*. (2017). Sequencing was conducted at the Illumina Sequencing Genomics Resources Core Facility at Weill Cornell Medical College with 1 x 50bp (SE) run on the HiSeq2500 platform. Raw sequencing data from maize and sorghum with the same experimental design and cold treatment were previously deposited at NCBI (http://www.ncbi.nlm.nih.gov/bioproject) under accession number PRJNA344653^7^. The raw reads were quality filtered, and adaptors were removed from the data with the sequence pre-processing tool Trimmomatic (v 0.38)^29^ (MINLEN = 36, LEADING = 3, TRAILING = 3, SLIDINGWINDOW= 4,15). The trimmed reads were mapped to the corresponding reference genome for each species using GSNAP^30^(v 2018-03-25)^29^ (-B 4 -N 1 -n 2 -Q -nofails format=sam). Genome assemblies of *Setaria italica* (v2.2)^31^, *Panicum virgatum* (v4.1)(DOE-JGI, http://phytozome.jgi.doe.gov/), *Zea mays* (APGv4)^32^, and *Sorghum bicolor* (v3.1.1)^33^ were downloaded from Phytozome version 12.1. Genome assemblies for *Panicum miliaceum* and *Pennisetum glaucum* were downloaded from NCBI34 and the Gigascience Database (http://dx.doi.org/10.5524/100192)^35^, respectively. Samtools (v 1.9)^36^ was used to convert the raw SAM output from GSNAP to sorted BAM files. FPKM (Fragments Per Kilobase of transcript per Million mapped reads) values were calculated using sorted bam files with cufflinks (v2.2)^37^. Genes were classified as expressed if their averaged FPKM values at all time points under both treatment and control conditions were ≥11^4^. HTSeq (v 0.6.1) was used to extract the number of reads in each RNA-seq library that were mapped to annotated exons of each gene in each species using union mode^38^. Read counts were used to identify cold-responsive genes by comparing the expression of genes in treatment vs. control samples, with differentially expressed genes defined as having adjusted p-value <0.05 and absolute log2 of fold change ≥ 2 at any of the six time points using DESeq2^39^. Nonresponsive genes were defined as those meeting the definition of expressed genes with absolute log2 of fold change of between treatment and control value ≤ 0.5 at all time points. Raw sequencing data that were not already public were deposited into NCBI’s SRA. See data availability statement.

### Quantifying Gene Features

Genomic features of foxtail millet, switchgrass, maize, and sorghum were scored using the corresponding gff annotation file and the mRNA transcript that was scored as primary for each individual gene model. For pearl millet and proso millet, instead, all annotated genes were scored due to the lack of primary transcripts information. Annotation of UTR sequences was inconsistent across species. In pearl millet and proso millet, UTR annotations were absent while in maize, sorghum, switchgrass, and foxtail millet, only a partial set of genes included UTR annotations. When UTRs were present, their median lengths were approximately 200 bp (5’UTR) and 350 bp (3’UTR). These lengths were standardized for all species. The frequencies of all individual nucleotides (4 features) and dinucleotides (16 features) were calculated for each of six regions: the CDS, intron, estimated 5’UTR, estimated 3’UTR, 1 Kb upstream of the 5’UTR starting site, and 1 Kb downstream of the (Figure 3). Overall, 120 features were scored for each gene. The code used to calculate these features has been deposited in Github (https://github.com/shanwai1234/GenomeFeature).

For maize, additional non-genomic sequence features were scored as detailed by Dai and coworkers^16^. Briefly, the epigenetic features included DNA methylation (quantified separately in the CG, CHG, and CHH contexts), three histone modifications (H3K4me3, H3K27me3, and H3K27ac), and open chromatin (quantified by ATAC-seq)^40^. Diversity and evolutionary features included GERP (genomic evolutionary rate profiling) scores^41^, PAV (Presence–Absence variations) frequency, orthologous gene in close relatives, synonymous mutation rate (Ks), non-synonymous mutation rate (Ka), Ka/Ks value, minor allele frequency (MAF) distributions, and SNP density features. MAF distributions and SNP density were calculated from the maize 282 association panel with data downloaded from Panzea (https://www.panzea.org/)^42^. Ka and Ks values for maize genes were calculated based on orthologous genes in maize, sorghum, and foxtail millet, and the resulting values were obtained from the previous work^43^. A syntenic gene list for *Z. mays, S. biocolor, S. italica, S. viridis, O. sativa, B. distachyon,* and *O. thomaeum* was downloaded from Figshare (http://dx.doi.org/10.6084/m9.figshare.3113488.v1)^7^. Any missing values in the non-sequence-based feature set for maize were imputed using the median value for that feature across all genes. Summaries of feature values for foxtail millet, pearl millet, proso millet, switchgrass, sorghum, and maize are provided in Supplementary Table S3 and S4.

### Binning of Cold-Responsive and Nonresponsive Genes

A binning method was used to reduce the bias of baseline gene expression and to balance the number of genes in the cold-responsive and nonresponsive datasets for supervised machine learning classification. The joint set of all cold-responsive and nonresponsive genes was sorted and segmented into 12 bins (dodeciles) based on average expression value. Within each dodecile, all genes of the less abundant class (either cold responsive or nonresponsive) were included as potential data points for training and testing, while the more abundant class was randomly subsampled to provide equal numbers of cold-responsive and nonresponsive genes within that particular dodecile.

### Gene-Family Guided Splitting of Training and Testing Data Sets

Protein sequences were clustered into families using the Markov Cluster (MCL) Algorithm as previously described^24,44^. Gene families were defined using OrthoMCL clustering with an inflation index of 1.5^45^. Pairwise similarity of the protein sequence encoded by the primary annotated transcript of each gene was quantified using the e-value reported by BLASTP^46^. Training/testing dataset splits were conducted at the gene-family level, meaning that different genes belonging to the same gene family were never present in both the training and testing dataset simultaneously to prevent the supervised machine learning models from learning family-specific features.

### Random Forest Training, Classification, and Evaluation

Feature datasets were pre-processed using the scale and center transformation methods of the “preProcess()” function in the R package caret^47^. The Random Forest classification model25 was employed as implemented in the R statistical programming language (v3.4.4)^48^. The random forest models were built with the class label vector as a factor type using 1001 trees (ntree=1001). Models were run with “importance” set to “T” to also calculated the “Mean Decrease in Accuracy” measure of importance for each feature.

Receiver Operating Characteristic (ROC) Curves for each model were generated using the R package ROCR^49^. The “prediction()” function from the ROCR package was used to evaluate the model; the “performance()” function was used with the arguments “tpr” and “fpr” to generate curves; and the argument “auc” was used to calculate the area under the curves. Classification performance was assessed by examining the area under the receiver operating characteristic curve (AUC) for the testing data set.

The control line for expression data in the ROC curve plots indicates the accuracy with which genes can be classified as cold responsive or nonresponsive based solely on average FPKM values across all tested time points and treatments. False positive rates, true positive rates, and AUC for expression control were calculated using the Scikit-Learn package^50^.

### Cross-Species Prediction

Prediction models trained using data from either one or several species were used to make predictions for data collected from the other single species. The same gene-family-guided splitting approach described above was used to ensure that an individual gene family was not present in the training data from one species and the testing data from another species. Testing data were balanced using the same binning method described above. For each species used as a target for cross-species prediction, the amount of testing data used was subsampled to produce a testing dataset including the same number of genes as the testing data used for intraspecies prediction with 5-fold cross validation.

### Identifying Syntenic Orthologs

Coding sequence data for primary transcripts of *S. italica* (v2.2)^31^ and *P. virgatum* (v4.1) (DOE-JGI, http://phytozome.jgi.doe.gov/) were retrieved from Phytozome version 12.1. Coding sequence data for *P. miliaceum* and *P. glaucum* were obtained from NCBI (BioProject number PRJNA431363)^34^ and the GigaScience Database (http://dx.doi.org/10.5524/100192)^35^, respectively. The software and corresponding settings used to identify syntenic orthologs were described previously7 with minor modifications. The parameter settings for LASTZ51 were as described7 except that a 75% sequence identity threshold was used for alignment. The QuotaAlign algorithm was used for further processing with -quota set to 1:1 for comparisons between *S. italica* and *P glaucum,* and 1:2 for comparisons between *S. italica* and *P miliaceum* or *P. virgatum* due to whole genome duplication in *P. miliaceum* and *P. virgatum*. Other parameters used for QuotaAlign and the subsequent polishing procedure were as described previously^7^. The syntenic orthologous pairs between *S. italica* and *S. bicolor* were downloaded from Figshare (http://dx.doi.org/10.6084/m9.figshare.3113488.v1)^7^. The Syntenic gene list generated among *S. bicolor, S. italica, P. glaucum, P. miliaceum,* and *P. virgatum* is shown in Supplementary Table S6.

### Phylogenetic Analysis of Species

A set of 7,064 gene groups were identified with syntenic ortholog representatives in sorghum, foxtail millet, pearl millet, both subgenomes of proso millet, and both subgenomes of switchgrass (7 total gene copies). Multiple sequence alignments for the annotated coding sequences for all seven genes within a group were generated using MAFFT (v7.149) with the parameter setting L-INS-i^52^. Poorly aligned regions after multiple sequence alignment were eliminated using Gblocks (v0.91b) with the following settings: minimum number of sequences for a conserved position: 9; minimum number of sequences for a flank position: 14; maximum number of contiguous nonconserved positions: 8; minimum length of a block: 10^53^. StarBEAST2 (v 0.15.5)^54^ implemented in BEAST 2.5.155 employing a Bayesian Markov Chain Monte Carlo framework was used to estimate both species trees and divergence dates. Due to the computational intensity of the analyses, an ensemble of 12 separate StarBEAST2 runs was employed, using different sets of 50 loci that were selected randomly (without replacement) from the alignments. Each StarBEAST2 run used analytical population size integration, the uncorrelated lognormal clock model, an HKY nucleotide substitution model with empirical frequencies, gamma category count of 4, and proportion invariant of 0.2. A calibrated yule model was used as a prior for tree topology using the previously estimated divergence time between foxtail millet and sorghum of 26 million years ago as a reference^31^, which was derived from the divergence time between rice and Panicodeae at approximately 50 million years ago^56^. Two independent runs of 40 million generations (sampled every 5000) were conducted in each analysis and combined with LogCombiner (v 2.5.1) with 20% burn-in for the species tree. Effective sampling sizes and MCMC convergence were examined using Tracer (v 1.7.1)^57^. A maximum clade credibility tree was compiled with TreeAnnotator (v 2.4.7) after discarding the initial 10% burn-in and the tree was visualized using FigTree (v1.4.4)^58^.

### Clustering and GO Enrichment Analyses

Clustering analysis was performed using cold-responsive genes from foxtail millet, pearl millet, switchgrass, and proso millet as a whole. The log2 fold values were normalized by row and analyzed using the R k-means function with 20 groups. Groups with similar expression patterns were further merged into 13 clusters, including 4 early transcriptional response clusters, 4 late transcriptional response clusters, 2 continually changing clusters, and 3 unclassified clusters. The cluster patterns are shown in heat maps and in graphical format. Gene Ontology (GO) annotations were downloaded from phytozome (v 12.1) for foxtail millet and switchgrass and from published papers for proso millet34 and pearl millet^35^. GO enrichment analyses of gene sets in each cluster were performed using GOATOOLS59 with all annotated genes in the genome as background. GO terms were considered significantly enriched if p-value < 0.05 after controlling for false discovery rate using the Benjamini–Hochberg procedure.

## Acknowledgments

This work was supported by award number 2016-67013-24613 from the USDA National Institute of Food and Agriculture to J.C.S. and R.L.R. and by the U.S. Department of Energy, Office of Science, Office of Biological and Environmental Research Program under award number DE-SC0020355.

## Author contributions statement

X. M. and Z.L. conducted the analyses. X.D. calculated epigenetic, diversity, and evolutionary features of the maize model. Y. Z. and D.W.N. conducted cold-stress treatments and constructed RNA-seq libraries. S.M. and X.M. conducted electrolyte leakage analysis. J.C.S. and R.L.R. conceived the experiments. X.M. and J.C.S. drafted the manuscript. All authors reviewed the manuscript.

## Additional information

Raw sequencing data for foxtail millet, proso millet, pearl millet, and switchgrass are available at NCBI under BioProject accession number PRJNA650146; supplementary tables are available at Figshare (https://doi.org/10.6084/m9.figshare.12756431); the code used to calculate the genomic and gene model features is available at Github (https://github.com/shanwai1234/GenomeFeature).

The authors state that they have no competing interests.

## Supplementary materials

**Figure S1.**
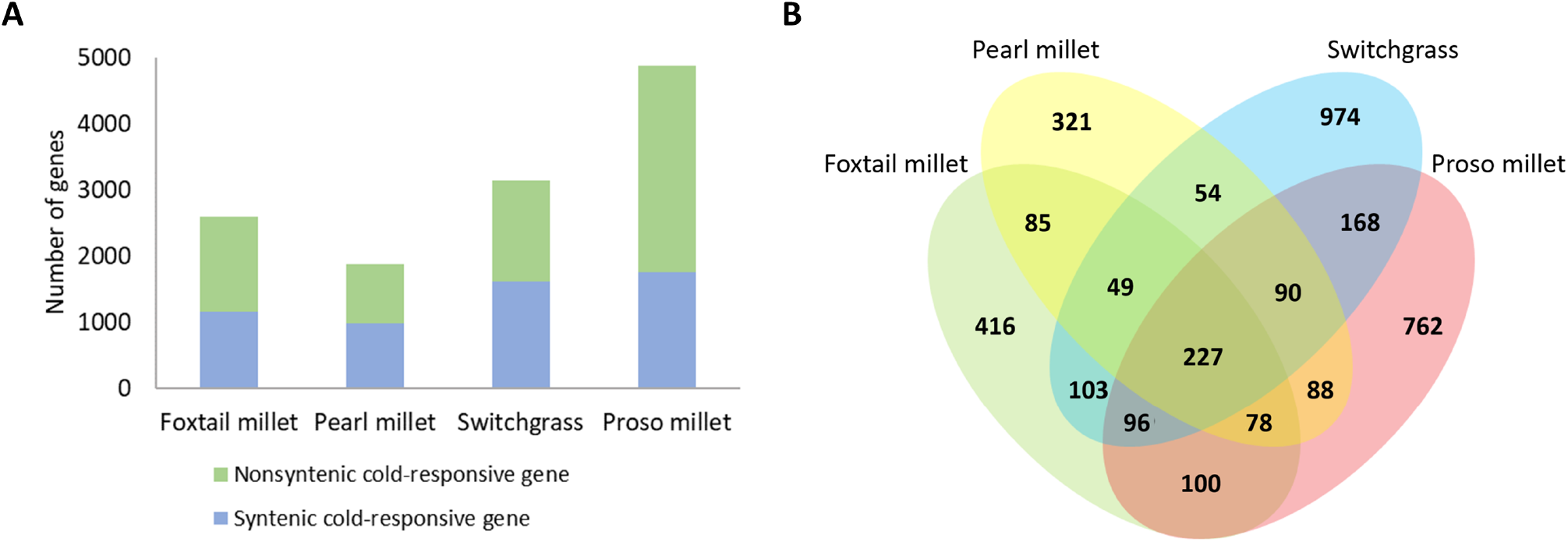
Conserved cold-responsive genes across foxtail millet, pearl millet, switchgrass, and proso millet. A. Proportions of syntenic orthologous genes among cold-responsive genes. B. Overlapping cold-responsive syntenic orthologs among the four species.

**Figure S2.**
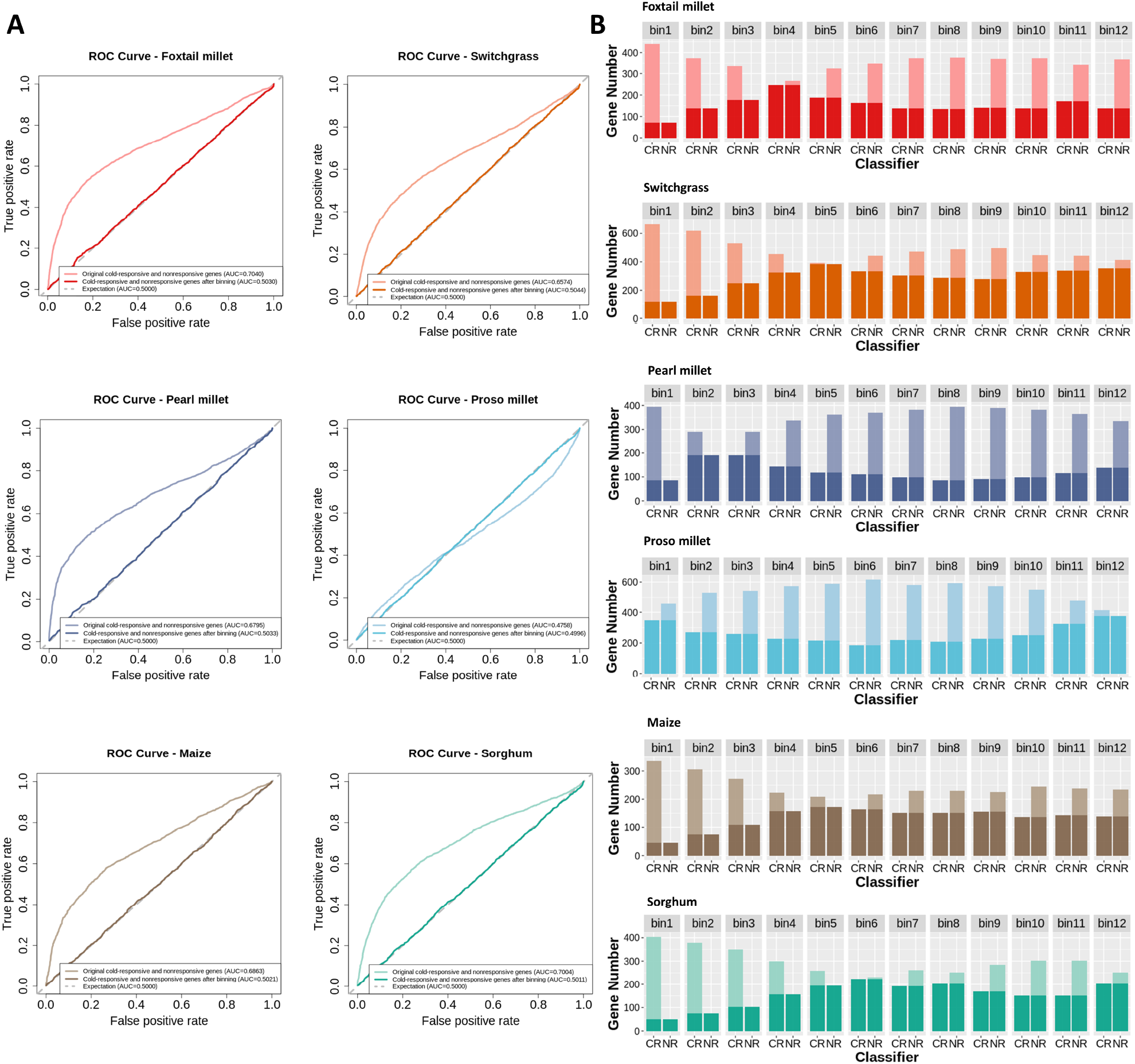
Baseline expression controls. A. Accuracy of genes being scored as cold-responsive genes solely based on average FPKM values before and after baseline expression control. B. Distribution of average FPKM values of cold-responsive genes (CR) and nonresponsive genes (NR), and training sets resampled from genes in dodeciles with balanced gene expression levels (darker color).

**Figure S3.**
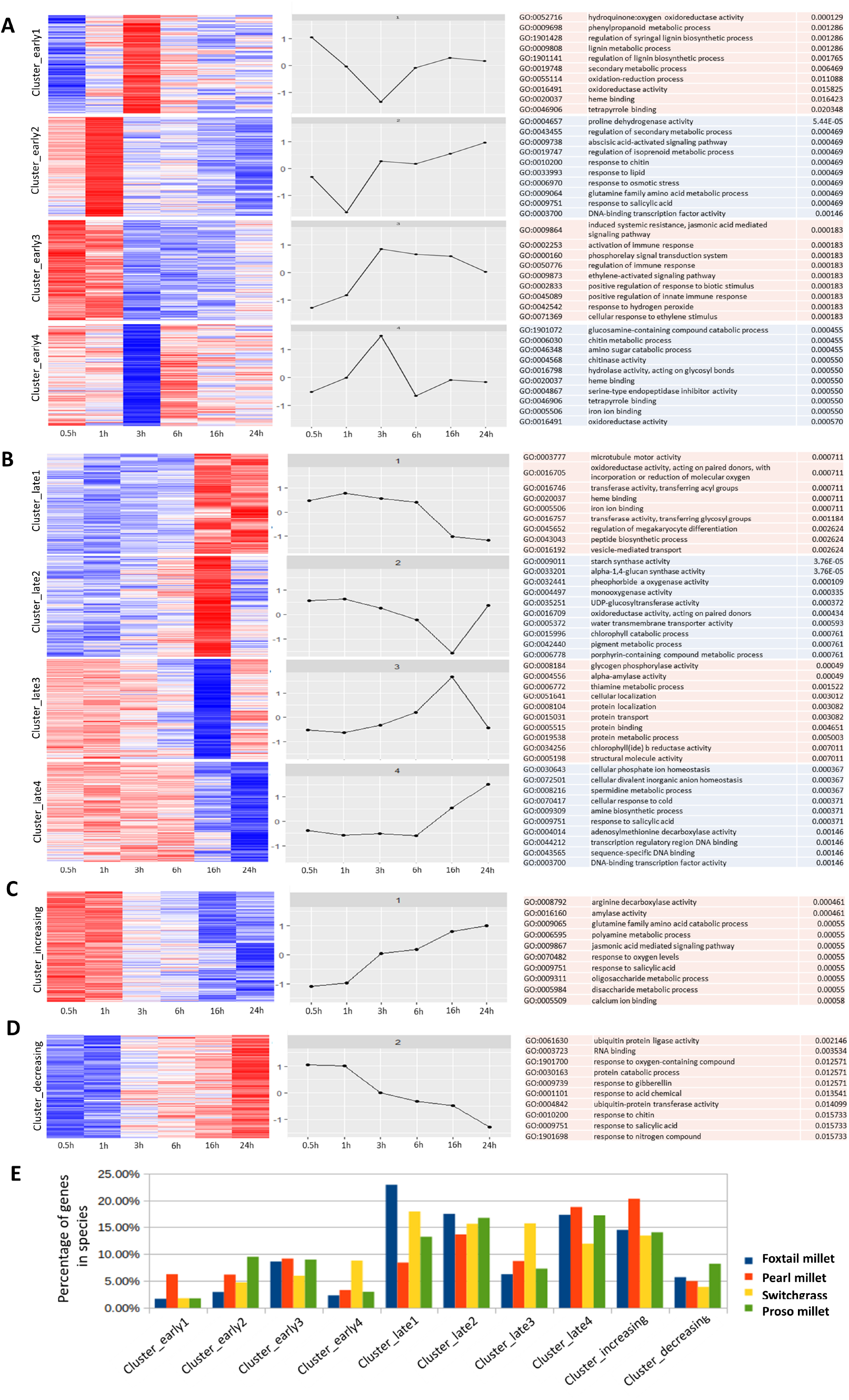
Gene expression clusters analysis. A-D. Cold-responsive genes from foxtail millet, pearl millet, switchgrass, and proso millet were analyzed using k-means clustering. This process identified eight major groups, as shown in heat map and graphical format, based on patterns of gene expression at different time points. (A) Clusters containing genes of early transcriptional responses to cold (30 min to 3 h); (B) clusters with late responded genes to cold (responded after 6h); (C and D) genes with continuously increasing or decreasing transcriptional levels within 24hrs. Enriched GO terms within clusters were shown in the last column. E. Percentage of genes of each of the four species distributed in clusters.

